# Replisome passage through the cohesin ring

**DOI:** 10.1101/2024.10.30.621121

**Authors:** Samson Glaser, Maxim I. Molodtsov, John F. X. Diffley, Frank Uhlmann

## Abstract

Following eukaryotic genome replication, the two newly synthesised sister chromatids remain paired by the ring-shaped cohesin complex, enabling their faithful segregation to daughter cells during cell divisions^1^. Cohesin topologically embraces DNA already before DNA replication^2–5^, and replisome passage through the cohesin ring is thought of as a fail-safe mechanism ensuring that cohesin entraps both replication products^6,7^. Whether replisomes indeed pass through cohesin rings remains unknown. Here, we use biochemical reconstitution^4,8,9^ and single molecule fluorescence microscopy to directly visualise replisome-cohesin encounters. We find that the translocating eukaryotic replicative Cdc45-Mcm2-7-GINS (CMG) helicase is frequently an obstacle to cohesin, but that the likelihood with which CMG passes cohesin increases in the presence of replisome components with known sister chromatid cohesion functions^7,10–14^, or by preventing cohesin from freely sliding along the template DNA. Cohesin retains topological DNA entrapment during CMG passage, suggesting that the helicase passes through the ring. The passage frequency increases further when fully reconstituted replisomes encounter cohesin rings, resulting in successful establishment of cohesion between the two replication products. Our findings demonstrate the existence of a simple mechanism that links genome replication with chromosome segregation, replisome passage through cohesin rings.

---

Sister chromatid cohesion provides the counterforce to mitotic spindle forces in the tug of war that aligns chromosomes on the cell equator^1^, in preparation for their faithful segregation when cohesin cleavage triggers anaphase. The ring-shaped cohesin complex is topologically loaded onto DNA already during G1 phase of the cell division cycle with help of its Scc2-Scc4 cohesin loader complex^4,15^. Following DNA replication, the same cohesin rings topologically embrace two DNAs^3,5,7,16^. How, during DNA replication, cohesin transitions from embracing one to embracing two DNAs remains unknown. A series of replisome components are known as cohesion establishment factors. In addition to their DNA replication roles, they aid successful sister chromatid cohesion establishment^7,10–14^. The genetic relationships between these factors^17,18^ suggest that more than one reaction co-ordinately contribute to cohesion establishment. Firstly, sister chromatids must be co-entrapped, and secondly replication-coupled cohesin acetylation stabilizes the resultant sister chromatid linkages^19–21^. A fail-safe mechanism to achieve the first requirement, co-entrapment of the two replication products, is replisome passage through cohesin rings. The cohesin ring diameter is ∼35 nm^6,22^, large enough in principle to allow passage of eukaryotic replisomes, which have a diameter of ∼25 nm^23^. On the other hand, DNA-bound proteins as small as 10 nm are known to form obstacles that cohesin struggles to overcome^24–26^. This small exclusion limit suggests that cohesin adopts a collapsed, or folded, conformational state when bound to DNA. Whether, therefore, the replisome can pass through cohesin rings remains unknown.

## CMG as a surmountable cohesin barrier

We began by visualising encounters of the replicative CMG helicase – the core component and main molecular motor of the eukaryotic replication fork – with cohesin. We purified fluorescently labelled budding yeast cohesin (tetramer complexes consisting of Smc1, Smc3, Scc1, Scc3) and its Scc2-Scc4 loader complex (Extended Data Fig. 1a). Fluorophore LD655-labelled cohesin exhibited DNA and Scc2-Scc4-stimulated ATPase activity, and could be loaded onto DNA in a salt-resistant manner, similarly to unlabelled cohesin (Extended Data Fig. 1b,c)^4,27^. Using a microfluidic flow cell, we loaded cohesin onto surface-tethered, stretched linear DNA. Following a high-salt wash to remove non-topologically bound cohesin, we visualised cohesin using total internal reflection fluorescence (TIRF) microscopy (Fig. 1a). Cohesin was seen as diffraction-limited spots moving along DNA in a diffusive manner. Photobleaching revealed that over 90% of these cohesin spots exhibited a single photobleaching step, indicating that most fluorescent foci correspond to single cohesin rings (Extended Data Fig. 1d). The diffusion coefficient increased with ionic strength in the imaging buffer. At 500 mM NaCl it reached *D* = 2.65 ± 0.21 µm^2^/s (mean ± S.E.M., n = 11, Extended Data Fig. 1e), comparable to diffusion coefficients reported for human and fission yeast cohesin^24–26^, or PCNA^28^. The value is substantially greater than observed for typical DNA-binding proteins that associate with DNA by electrostatic interactions^29^, consistent with loose topological entrapment of the DNA by these protein rings. Topological entrapment was additionally confirmed by DNA cleavage with a restriction enzyme, resulting in cohesin sliding off the free DNA end (Extended Data Fig. 1f).

**Figure 1.**
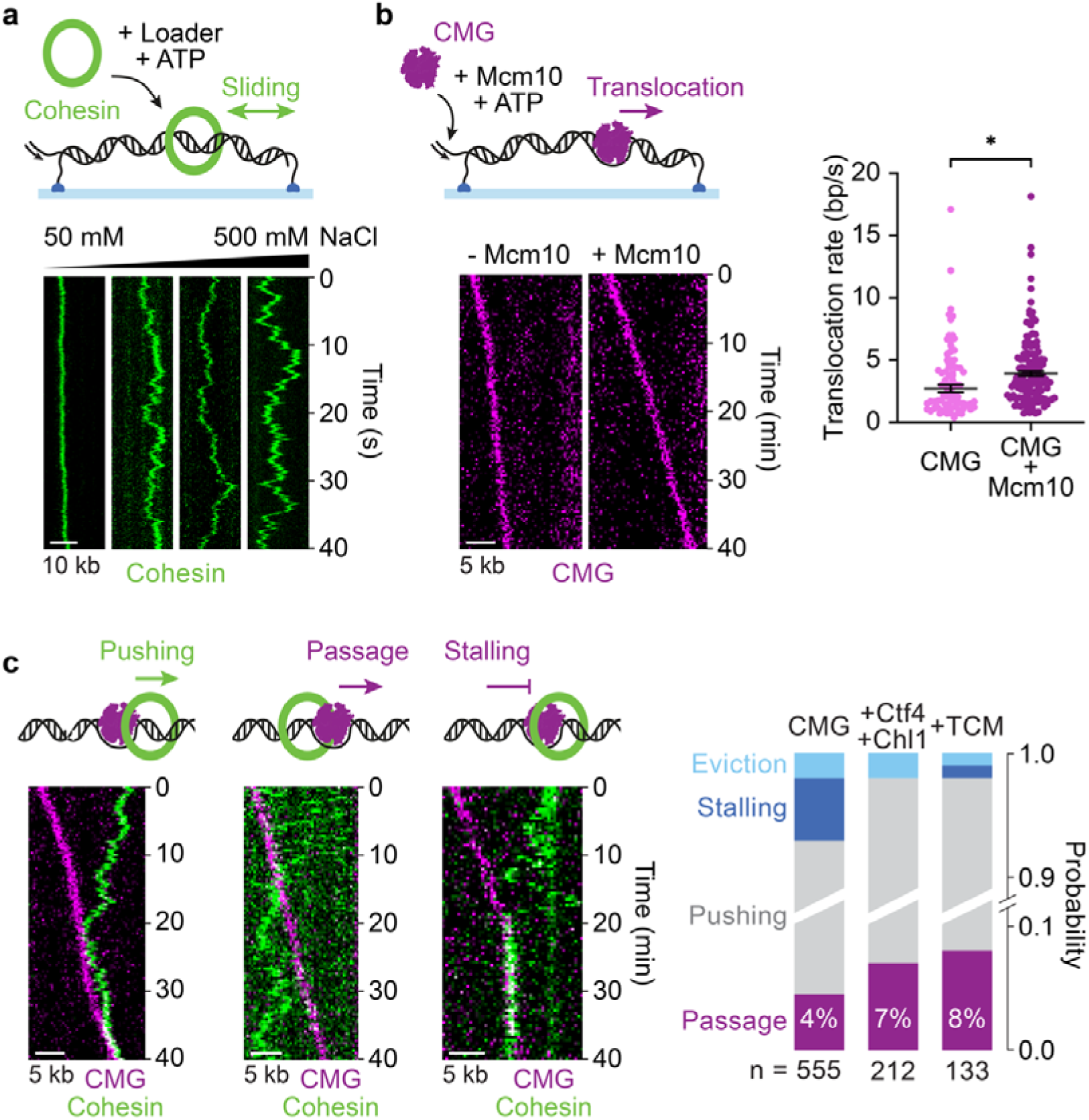
The CMG helicase is a surmountable cohesin barrier. **a,** Schematic of cohesin loading and sliding, as well as representative kymographs of cohesin loaded onto tethered DNA in a flow cell and imaged at the indicated salt concentrations. **b,** Schematic of CMG loading and translocation. CMG is loaded at a low ATP concentration, translocation is initiated by increasing ATP, with or without Mcm10 addition. Representative kymographs of CMG translocation in the presence or absence of Mcm10. Distributions of segment translocation rates for CMG without and with Mcm10 are shown. Black lines and error bars represent the weighted mean and standard error (* *p* < 0.001, unpaired t-test). **c,** Schematics and representative kymographs from observed outcomes of CMG-cohesin encounters. Frequencies were aggregated from four biological replicates with CMG and two biological replicates with CMG+Ctf4+Chl1 and CMG+TCM, respectively. n is the total number of observed encounters (*p*(CMG vs. CMG+Ctf4+Chl1) = 0.0163, *p*(CMG vs. CMG+TCM) = 0.0865, unpaired t-test).

Next, we purified and fluorescently labelled budding yeast CMG and observed equally efficient bulk DNA unwinding by both labelled and unlabelled CMG (Extended Data Fig. 2a,b). To visualise CMG translocation at the single-molecule level, we tethered a linear forked DNA with a free 3’ overhang to the flow cell surface, followed by helicase binding in presence of a low ATP concentration, and then helicase activation by increasing ATP^30^. This protocol resulted in fluorescently labelled CMG moving unidirectionally along the DNA (Fig. 1b). As previously seen^31^, addition of the CMG cofactor Mcm10 resulted in an increased translocation rate (Extended Data Fig. 2c). In this experimental setup, unwound DNA reanneals behind the translocating CMG. To confirm that CMG unwinds DNA, we included the single-strand binding protein RPA and the DNA dye SYTOX Orange, which preferentially binds double stranded DNA. Over 80% of translocation events were accompanied by the loss of the SYTOX Orange signal, confirming that the CMG helicase unwinds DNA during translocation (Extended Data Fig. 2d). In the following, we performed experiments without RPA, resulting in CMG movement in the 3’-5’ direction along the forked strand, which then reannealed.

To visualise cohesin-CMG encounters, we first loaded cohesin onto the forked DNA, and then bound and activated the CMG helicase. When a translocating CMG encountered cohesin, we observed one of four outcomes (Fig. 1c). In most cases, CMG appeared to push cohesin or to confine its available space for diffusion along the DNA (‘pushing’). However, in a small fraction of instances (24 out of 555 events), cohesin was passed by CMG upon encounter, and cohesin continued diffusive motion along DNA on the other side (‘passage’). Other low frequency events were apparent cohesin eviction (‘eviction’), or CMG stalling upon encounter (‘stalling’). While the passage frequency was low, we note that passage was never previously observed across an approximately similarly-sized quantum dot (CMG 18.9 nm, quantum dot 19.5 nm)^24,32^.

To explore the physiological relevance of cohesin passage events, we investigated whether cohesion establishment factors that associate with CMG (Extended Data Fig. 2a) alter the outcome of CMG-cohesin encounters. Addition of Ctf4 and Chl1, or of Tof1-Csm3 and Mrc1 (the TCM complex) increased the frequency of passage events, in both cases by around two-fold (Fig. 1c). By binding CMG, these proteins increase the overall size of the translocating apparatus^32–34^. However, instead of augmenting CMG barrier function, these cohesion establishment factors facilitated cohesin passage. Chl1 and TCM make direct physical contact with cohesin^34–36^, and these contacts likely engaged cohesin in ways that facilitated passage.

## Converging CMG helicases pass cohesin

Topologically loaded cohesin is free to slide along bare DNA that is tethered to our flow cell surface. Cohesin sliding is likely much more restricted on physiological chromatin substrates due to the presence of abundant nucleosomes^24–26^. To investigate the effect of constrained cohesin mobility, we took two approaches. In the first approach, we designed a DNA template featuring 3’ forked overhangs at both ends, allowing the loading of two CMG helicases that approach each other in 3’-5’ direction on opposite DNA strands. Upon encounter, the two CMGs either stalled (57% of events) or passed each other (43% of events) (Fig. 2a).

**Figure 2:**
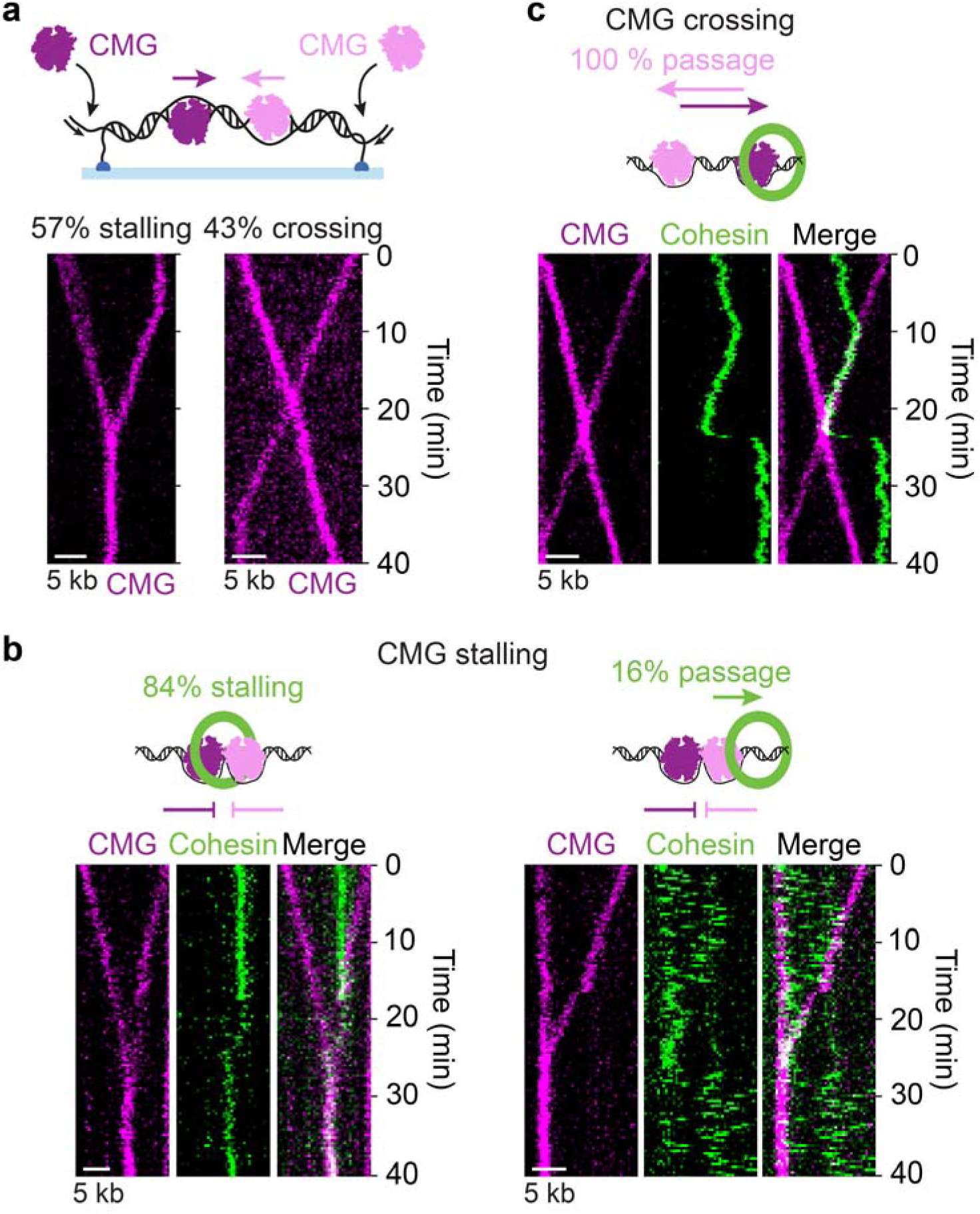
Cohesin passage by converging CMG helicases. **a,** Schematic and representative kymographs of converging CMG helicases, which either stall (left) or cross each other (right) upon encounter (n = 87). **b,** Schematics and representative kymographs of cohesin between two stalling CMGs. A cohesin stalling (left) and a cohesin passage (right) event are shown (n = 25). **c,** Schematic and representative kymograph of cohesin between two crossing CMGs. One of the CMGs passes cohesin in all cases (n=12).

Next, we investigated the fate of cohesin between two converging CMGs. As CMGs approached each other, the range of cohesin movements was typically constrained up to the point where CMGs met. In cases when CMG movement stalled, cohesin became trapped (Fig. 2b, left), but in 16% of cases cohesin passed one of the two CMGs (Fig 2b, right). This passage rate is substantially greater than the rate that we observed when freely mobile cohesin encountered one CMG. The observation suggests that an increased contact time of cohesin with CMG facilitates passage.

In all those cases where two converging CMG helicases crossed one another, cohesin was pushed by one CMG over the other (Fig. 2c). After passing one of the CMGs cohesin resumed diffusive motion along the DNA, suggesting that it retained topological DNA embrace during passage. We did not observe cohesin eviction during CMG-CMG passage events, hinting at the possibility that cohesin remained topologically engaged with DNA throughout the passage event. From these observations, we can conclude that CMG readily passes cohesin rings, if these are prevented from sliding away. The previously observed collapsed conformational state of DNA-bound cohesin, which precludes passage of many DNA-bound obstacles^24–26^, was therefore unfolded by an advancing CMG helicase, with help of cohesion establishment factors.

## CMG passage of immobilised cohesin

We used a second experimental approach to study the mechanics of CMG-cohesin encounters. Following loading onto DNA, we tethered cohesin to the flow cell surface by means of a V5 epitope-specific antibody, recognising a V5 epitope tag fused to Smc3, that we functionalised with a 20 kDa (∼150 nm) PEG-biotin linker (Extended Data Fig. 3a). Introduction of this antibody into the flow cell resulted in immobilisation of previously diffusive cohesin (Fig. 3a). An unspecific functionalised antibody, introduced as a control, or a V5 antibody lacking PEG-biotin (Extended Data Fig. 3b), did not overtly affect cohesin mobility.

**Figure 3:**
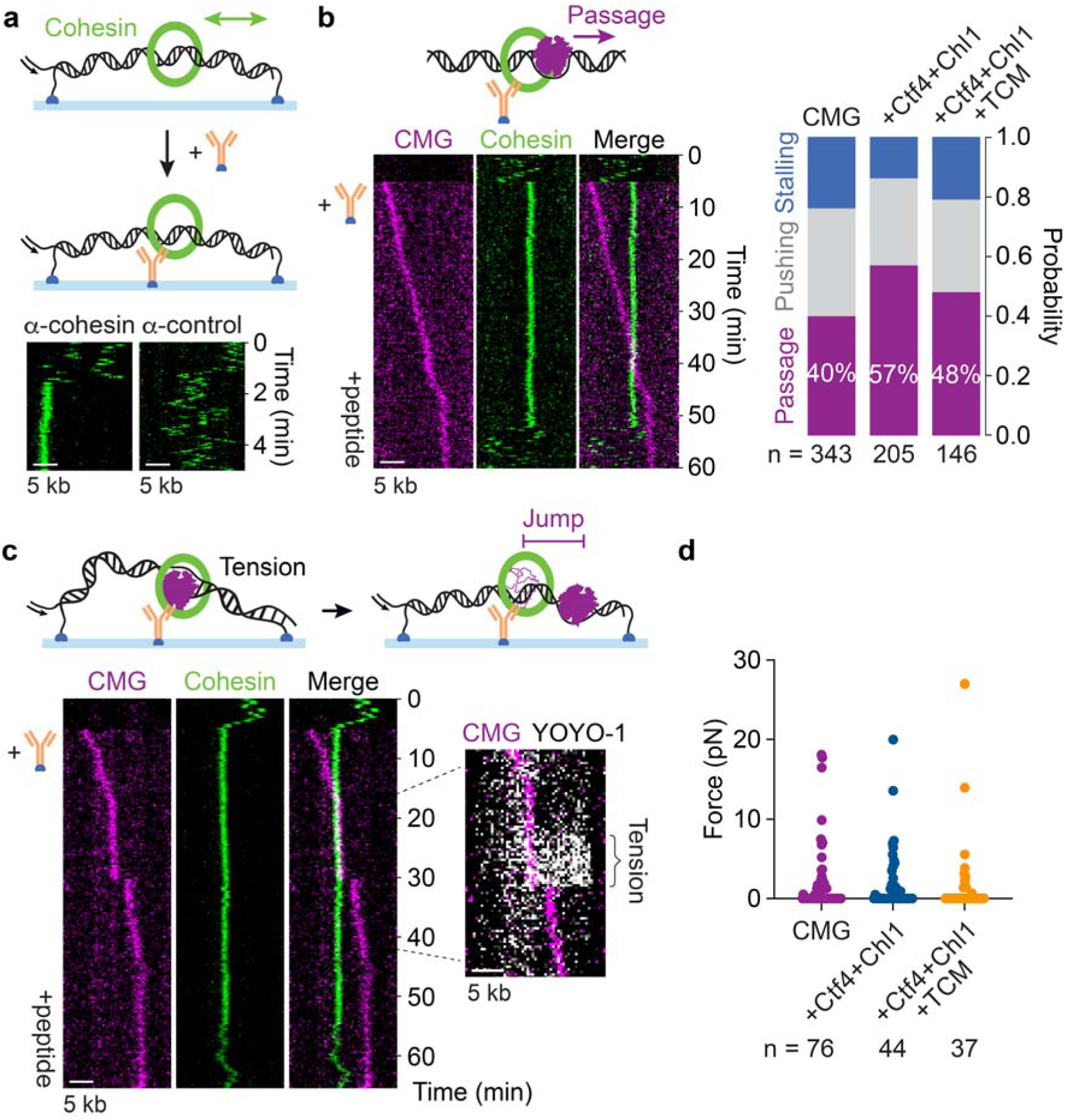
CMG passes immobilised cohesin. **a,** Schematic of cohesin immobilisation using a functionalised antibody. Representative kymographs of DNA-bound cohesin after adding a biotin-coupled cohesin-specific (left) or unspecific (right) antibody. **b,** Schematic and representative kymograph of CMG passing immobilised cohesin. A diffusive cohesin was imaged for 5 min. After antibody immobilisation and CMG binding and activation, cohesin and CMG were imaged for 40 min, before cohesin was mobilised again by peptide elution. Outcomes from three biological replicates experiments were scored and aggregated, as well as two replicates with CMG+Ctf4+Chl1 and four replicates with CMG+TCM, respectively. n is the total number of observed encounters (*p*(CMG vs. CMG+Ctf4+Chl1) = 0.093, *p*(CMG vs. CMG+TCM) = 0.460, unpaired t-test). **c,** Schematic and representative kymograph of CMG stalling and jumping on encountering immobilised cohesin. Continued CMG translocation during stalling results in DNA stretching, evident by increased YOYO-1 staining in front of CMG. **d,** Forces generated by CMG leading up to a jump, calculated from the DNA stretch factor and jump size. The force is assumed to be negligible if no jump was observed. n is the total number of passage events.

We then performed CMG-cohesin encounter experiments with immobilised cohesin. We started the experiment by imaging diffusive cohesin, to confirm its topological binding to DNA, and then introduced the V5 antibody to immobilise cohesin. Next, we bound and activated CMG. Cohesin passage now became the most frequent outcome, with 40% of CMG-cohesin encounters resulting in passage (Fig. 3b). Other outcomes were cohesin pushing, presumably following cohesin detachment from the antibody, and CMG stalling (Extended Data Fig. 3c). Inclusion of Ctf4 and Chl1, or of TCM, resulted in a further small increase in passage frequency. Following passage, we introduced V5 peptide into the flow cell, which resulted in cohesin elution from the antibody tether. Cohesin now resumed diffusive motion along the DNA, indicating that it retained topological DNA association following CMG passage (Fig. 3b and Extended Data Fig. 3d). These observations confirm that CMG passage is greatly facilitated by restricting cohesin’s ability to escape from the advancing helicase.

### Force build-up during CMG-cohesin encounters

When we examined CMG traces while passing immobilised cohesin more closely, we noticed transient stalling, followed by small jumps, in approximately half of the encounters (Fig. 3c and Extended Data Fig. 3e). No interruptions to CMG progression were apparent in the other half of passage events. CMG stalls and jumps might result from cohesin forming a temporary obstacle, holding up CMG, while the helicase continues to reel in DNA. Continuing translocation along DNA would lead to tension build-up, resulting in a jump once the tension is released upon passage. To explore this scenario, we conducted three-colour TIRF experiments in which we simultaneously visualised CMG, cohesin, and the DNA stain YOYO-1. DNA tension facilitates dye intercalation, so that increased YOYO-1 staining can serve as an indicator for DNA tension^37^. The YOYO-1signal ahead of, but not behind, CMG indeed increased during stalling and dropped again coincident with the jump (Fig. 3c). This observation is consistent with tension build-up during stalling and release upon cohesin passage. Following these passage events and V5 peptide elution, cohesin again resumed diffusive motion along the DNA, confirming that cohesin retained topological DNA embrace also following these force-coupled passage events.

We next wanted to know how much tension force is generated by CMG before a jump. Might the accumulated force have broken the cohesin ring? From the measured change in DNA segment length in front of the CMG helicase before and after the jump, and the known DNA force-extension relationship^38^, we derived the stretching force that preceded individual CMG jump events (Fig. 3d, see the Methods for details). This analysis revealed that the vast majority of passage events occurred at well below 20 pN, and therefore well below the force required to rupture the cohesin ring at its weakest interface^5^. The most parsimonious explanation for these observations is a mechanical model in which CMG passes through the cohesin ring, but in which passage is sometimes obstructed by cohesin’s collapsed or folded conformational state^24^. A passage-competent cohesin shape might be reached in a conformational search, aided by cohesion establishment factors that interact with cohesin, as well as by force build-up by the advancing CMG helicase.

### CMG passage through the cohesin ring

The above experiments have established that cohesin remains topologically bound to DNA following CMG passage. However, what happens to cohesin at the moment of passage remained unresolved by these observations. Does CMG indeed pass through the intact cohesin ring, or must the ring temporarily open by disengagement of one of its interfaces? To differentiate between these scenarios, we challenged the cohesin-DNA interaction during CMG passage in side-flow experiments. In these experiments, cohesin was loaded onto DNA and we then applied a strong side flow while tethering cohesin to the flow cell surface using the V5 PEG-biotin antibody (Fig. 4a). As a result, DNA was held by cohesin in an extended V-shape after we stopped the side flow. Cohesin elution from the antibody tether by V5 peptide addition released the DNA, which regained a linear geometry while cohesin resumed diffusive motion. These observations confirm that a topologically loaded cohesin resisted a perpendicular pull on the DNA. In around 10% of observed cases, cohesin spontaneously released the DNA while itself remaining surface-bound (Extended Data Fig. 4).

**Figure 4:**
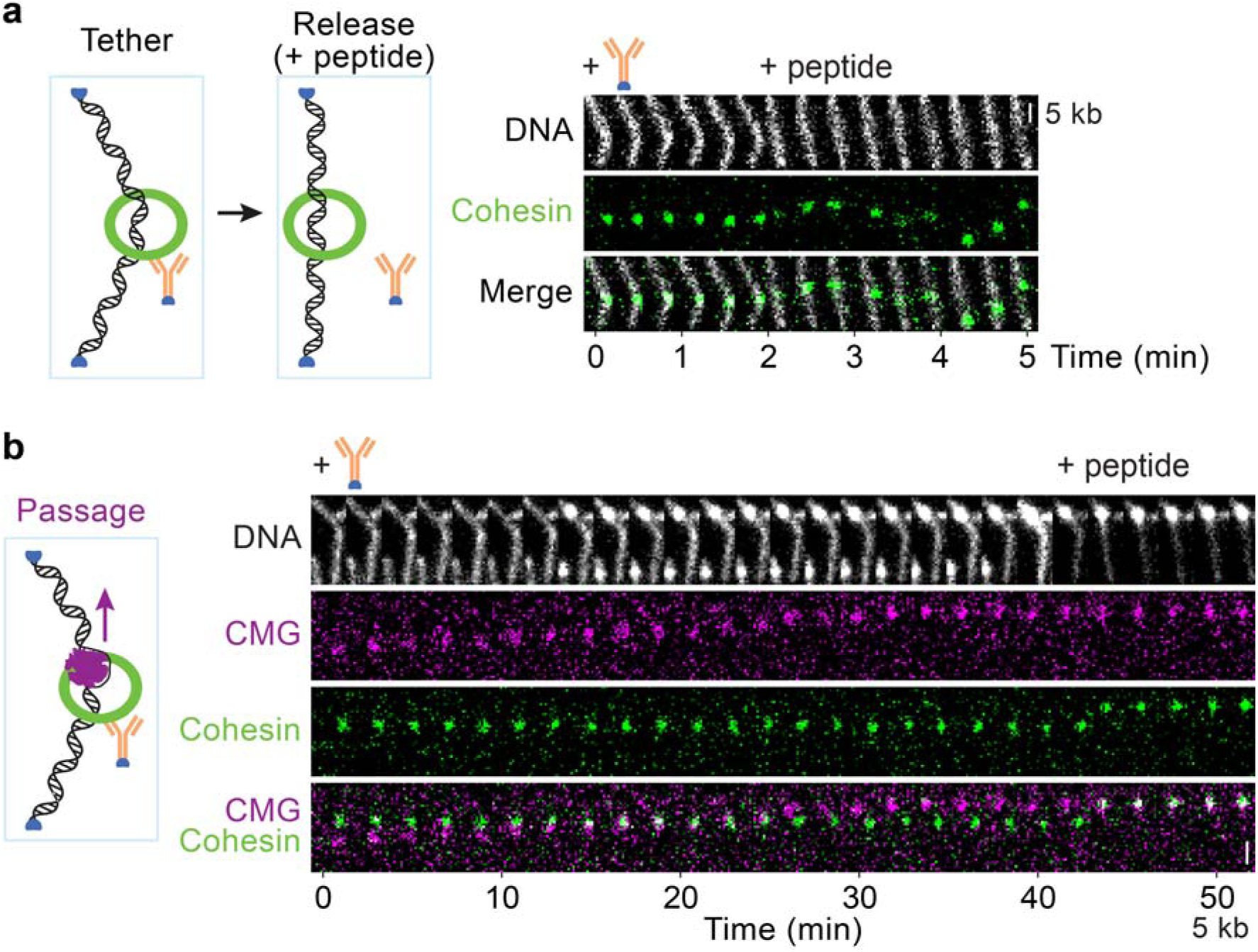
Cohesin retains DNA association during CMG passage. **a,** Schematic and example time series of side flow-stretched DNA, tethered by cohesin (Tether). Upon peptide addition, DNA and cohesin are released from the antibody (Release). **b,** Schematic and representative time series of CMG passage along flow-stretched DNA and past immobilised cohesin. Cohesin remains DNA bound during passage, as well as after peptide elution.

If cohesin transiently opened during CMG passage, the applied perpendicular pull should result in DNA release and return to its linear conformation. In contrast to this expectation, DNA retained its extended V-shaped geometry during all observed CMG passage events, indicating that DNA remained topologically entrapped by the cohesin ring while CMG traversed through its centre (n=4, Fig. 4b). Only peptide elution of cohesin from the antibody tether after CMG passage eventually allowed DNA recoil, while cohesin remained DNA-bound. These observations suggest that CMG passed through an intact cohesin ring. The alternative explanation, that the cohesin ring opened but DNA remained cohesin-bound by electrostatic interactions, is made unlikely by the previous observation of spontaneous detachment events, during which electrostatic cohesin interactions were visibly insufficient to retain DNA (Extended Data Fig. 4). Furthermore, there are no known direct CMG-cohesin interactions^35^ that could have maintained contact with an open cohesin ring.

### Establishment of sister chromatid cohesion

Finally, we visualised fully reconstituted replisomes in the act of replicating our forked DNA substrate^30^ and encountering cohesin. We bound CMG to the primed fork in the presence of the non-hydrolysable ATP analogue adenylyl-imidodiphosphate (AMP-PNP) to prevent any onset of DNA unwinding. We then added the remaining replisome components Mcm10, Ctf4, Tof1-Csm3, Mrc1, Pol α, Pol δ, Pol ε, RFC, PCNA, and RPA (Extended Data Fig. 5a), and exchanged AMP-PNP for NTPs and dNTPs. DNA replication was visualised using SYTOX Orange, which stains the leading strand DNA product as a bright spot of increasing intensity that moves along the template DNA (Fig. 5a). The lagging strand product remains tethered to the surface such that the DNA stain behind the replisome remains unchanged from that of the template DNA. By tracking the centres of leading strand foci we quantified the DNA synthesis rate, revealing a range of rates with an average of 19.9 ± 0.6 bp/s (mean ± standard error, Extended Data Fig. 5b), consistent with previous observations^9,30^. Omitting TCM from the experiment, as expected^9,30^, resulted in substantially lower replication rates, confirming that these cohesion establishment factors are proficient in fulfilling their replication roles in our assay.

**Figure 5:**
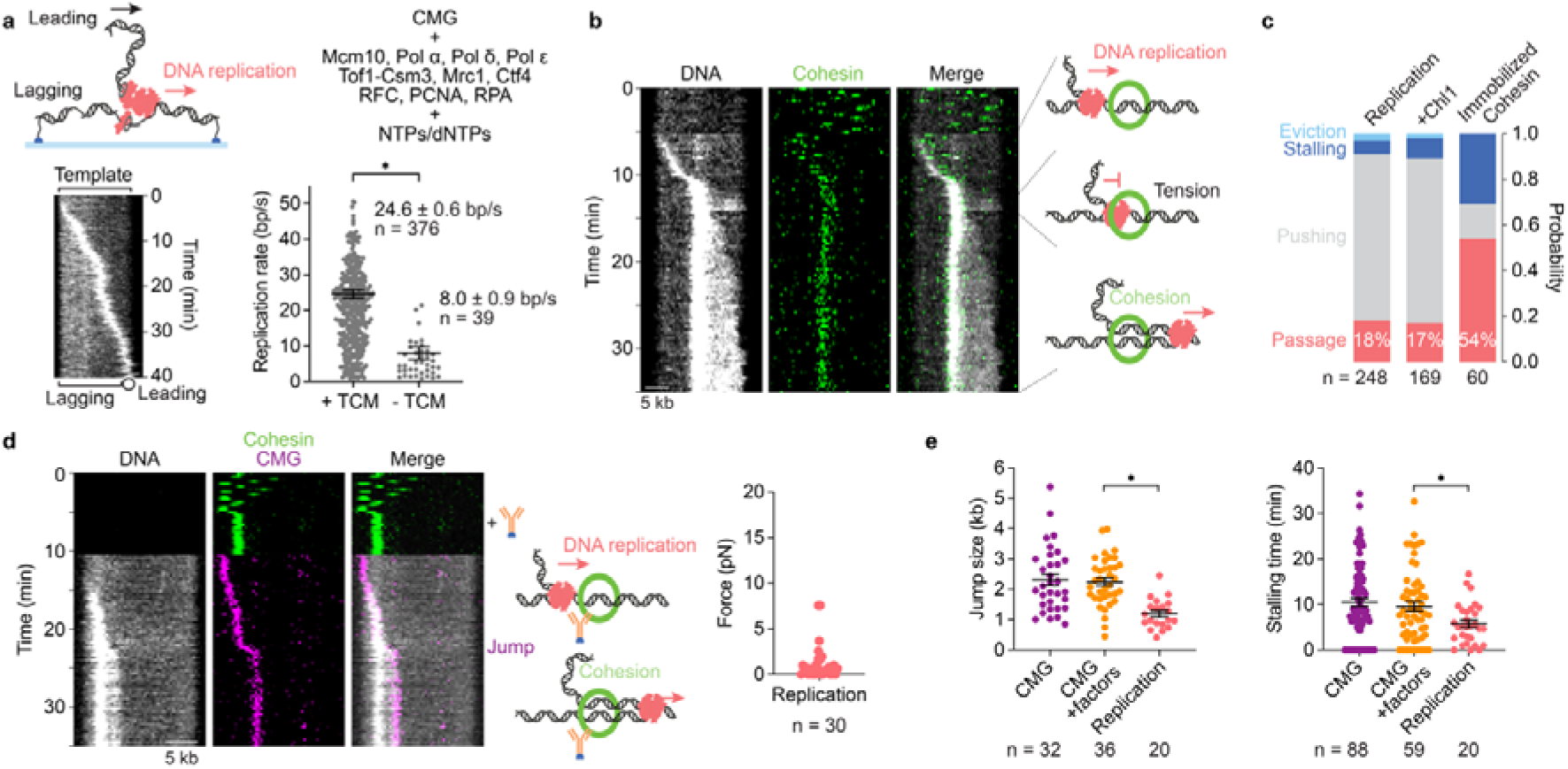
Replisome passage through the cohesin ring. **a,** Schematic and example kymograph of the DNA replication assay. The DNA template, lagging strand replication product, as well as the globular-coiled leading strand product are indicated. Intensity fluctuations of the latter are likely due to diffusive motion in and out of the TIRF excitation volume. Replication rates in the presence and absence of TCM are shown. Each data point represents a DNA replication segment. Black lines and error bars represent the weighted mean and standard error (* *p* < 0.0001, unpaired t-test). n is the total number of segments. **b,** Schematics and representative kymograph of the events during replisome passage through the cohesin ring. First, the replisome approaches cohesin. On encounter, a stretch force transiently builds up (distributed SYTOX Orange intensity increase). Following passage, progressive SYTOX Orange intensity increase results from the two newly synthesised leading and laggings strands. **c,** The outcomes of replisome-cohesin encounters were aggregated from 4 biological replicates (Replication), 2 replicates (+ Chl1) and 2 replicates (immobilised cohesin), n is the total number of observed encounters. **d,** Example kymograph of cohesin encounters by replisomes including LD655-labelled CMG. Only cohesin was imaged until 10 min, then excitation was changed to image CMG and DNA. Forces generated in the run-up to jumps were calculated from the DNA stretch factor and jump size. The force is assumed to be negligible if no jump was observed. n is the total number of passage events. **d,** Comparison of jump sizes and stalling times during CMG-, CMG with cohesion establishment factor-, and replisome-cohesin encounters. Black lines and error bars represent means and standard errors (* *p* < 0.0001 jump size, *p* = 0.0038 stalling time, unpaired t-tests).

To visualise replisome-cohesin encounters, we loaded cohesin onto the template DNA before replisome assembly and replication initiation. Upon replisome encounters with cohesin we observed two main outcomes. In many cases, the advancing replication machinery appeared to push cohesin (Extended Data Fig. 5c). Notably, however, 18% of encounters resulted in cohesin passage by the replisome. Replisome passage in turn occurred in two ways, with establishment of sister chromatid cohesion, or without (Fig. 5b, and Extended Data Fig. 5d, discussed further below). Despite the large size of a complete replisome, the cohesin passage frequency was substantially greater when compared to CMG-only cohesin encounters. Replisome stalling or cohesin eviction upon encounter remained rare events.

The increased replisome passage probability, as compared to CMG-only cohesin encounters, suggests that replisome components beyond the known cohesion establishment factors promote replisome-cohesin passage. These components might include Pol ε, whose truncation causes sister chromatid cohesion defects^39^. On the other hand, the cohesion establishment factor Chl1 did not increase the passage frequency (Fig. 5c). This finding could be explained by a recently suggested role of Chl1 in promoting topological cohesin loading ahead of the replisome^36^, a function that would be obscured in our experimental setup where we complete topologically loading of cohesin onto DNA before initiation of DNA replication.

Most observed instances where the replisome passed cohesin resulted in the establishment of sister chromatid cohesion (55 out of 71 passage events). Cohesion establishment was manifest by the following observations. The leading strand DNA that was already synthesised at the time of the cohesin encounter remained tethered by cohesin, seen as a mass of constant SYTOX Orange staining intensity that colocalised with cohesin for the remainder of the experiment (Fig. 5b). The DNA intensity ahead of cohesin, in turn, progressively increased as the replisome continued DNA synthesis, indicative of juxtaposition of both the newly synthesised leading and lagging strands. On the other hand, a minority of cohesin encounters did not result in cohesion establishment and the leading strand product continued traveling along the template DNA with the advancing replisome. A possible explanation for passage events without cohesion establishment is absence of a mechanism that prevents the leading strand product from sliding through and out of the cohesin ring (Extended Data Fig. 5d). These results are to our knowledge the first live observations of sister chromatid cohesion establishment while a replisome passes a cohesin ring.

### Replisome passage through the cohesin ring

When we inspected DNA traces during sister chromatid cohesion establishment more closely, we saw in around a third of passage events (23 out of 71) that the SYTOX Orange intensity ahead of cohesin transiently increased upon replisome-cohesin encounter, before the replisome passed cohesin (Fig. 5b). This observation suggests that tension sometimes builds up during encounters, before being released upon cohesin passage.

To quantify how much force the replisome exerts on cohesin before passage in these cases, we turned to replisome encounters with immobilised cohesin, tethered to the flow cell surface using the V5 PEG-biotin antibody. During encounters with immobilised cohesin, the replisome passage frequency increased to over half of all events, with successful cohesion establishment in 23 out of 32 passage events (Fig. 5c and Extended Data Fig. 6). We next repeated these cohesion establishment experiments with immobilised cohesin, but using replisomes containing fluorophore-labelled CMG. This setup allowed us to measure replisome jump sizes upon cohesin encounter (Fig. 5d). DNA tension forces that we derived from the jump sizes, as in the case of CMG-cohesin encounters, remained well below 20 pN, suggesting that cohesin rings were not broken by passing replisomes. Furthermore, jump sizes were shorter and stall times briefer during replisome-cohesin encounters, when compared to CMG-cohesin encounters (Fig. 5d). Taken together, our results suggest that the replisome passed through cohesin rings to leave the two replication products trapped inside. While the protein components of the replisome facilitate passage, as evident by less frequent and shorter jumps, the mechanical requirement of bringing the cohesin ring into a passage-competent conformation remains detectable as occasional transient replisome stalling events.

## Discussion

In this study, we visualise how a biochemically reconstituted eukaryotic replisome encounters cohesin rings that topologically embrace the template DNA. These experiments illuminate a key moment during the replication and propagation of eukaryotic genomes, the time when the two newly synthesised sister genomes are embraced by cohesin. Previous studies have highlighted the ease with which cohesin’s topological interaction allows it to slide along the DNA, but also cohesin’s difficulty to overcome small obstacles^24–26^. It is therefore thought that cohesin exists in a collapsed conformational state when bound to DNA. On the other hand, much of the cohesin ring circumference is made up of flexible stretches of coiled coil without a fixed conformation^22^. High speed atomic force microscopic visualisation of the related condensin complex further exemplifies the vast conformational space that SMC complexes sample^40^. We envision that cohesin mostly exists as a flexible and malleable ring on DNA. If approached from one side, even by an object smaller than its nominal diameter, a collapsed or folded ring will be displaced. On the other hand, if the ring is prevented from escape, an approaching structure will unfold the ring, enabling passage by objects of up to ring-diameter size. These considerations explain why preventing cohesin ring sliding substantially increases the likelihood of replisome passage and of sister chromatid cohesion establishment. Cohesin’s diffusive motion is greatly restricted by nucleosomes^24–26^ and circumstantial evidence suggests that nucleosomes influence the outcome of replisome-cohesin encounters. In crude *Xenopus* egg extracts, using a bare DNA substrate, replisome-cohesin encounters mostly led to cohesin pushing^41^, while similar experiments using chromatinised DNA resulted in replisome passage as the most frequent outcome^26^. Chromatin immunoprecipitation analyses in budding yeast suggested that cohesin does not change its position while a replisome passes^7^. How nucleosomes affect the outcome of replisome-cohesin encounters, and the detailed observations of what happens when replisomes meet cohesin *in vivo*, remain important areas for future studies.

Cohesion establishment factors increase the frequency with which the replisome passes, rather than pushes, cohesin rings. While increasing the overall size and overt steric hindrance of the replisome, we find that cohesion establishment factors aid cohesin passage. Amongst these factors, Tof1 is found at the very front of the replisome^32^, making Tof1 likely the first protein to encounter and contact cohesin^35^. The Tof1-cohesin interaction might serve two roles. On the one hand, engaging cohesin increases the time during which cohesin can sample its conformational space in search for a passage-competent shape. At the same time, Tof1 interactions might directly contribute to placing cohesin in a favourable orientation. Additional documented replisome cohesin interactions by Mrc1 and Chl1^35,36^, together with potentially further, as yet uncharacterised interactions, might chaperone cohesin along the replisome, similar to how multiple replisome-histone contacts guide histone inheritance during chromatin replication^42^. Chl1 promoted CMG helicase passage of cohesin, a positive effect that was no longer detectable during replisome-cohesin encounters. Protein interactions by other replisome components might have compensated for Chl1 in the latter experiments, while Chl1’s dominant role in the replisome context might be promoting topological cohesin loading ahead of the fork^36^.

Our experiments illustrate replisome passage through the cohesin ring, and co-entrapment of the two replication products. However, they do not fully establish what happens to cohesin at the moment of passage. Several aspects of our observations are easiest explained by replisome passage through cohesin rings that remain mechanically intact. DNA stretch forces sometimes build up during passage, as expected during a mechanical passage event, but not if a biochemical reaction was triggered by replisome-cohesin encounters. Nevertheless, we cannot exclude with certainty that replisome components catalyse the transient opening of a cohesin interface, without giving DNA a chance to escape. In the future, it will be important to establish whether the cohesin ATPase is engaged at the moment of replisome passage. Experiments to address this question are made difficult by the fact that the ATPase is essential for topological cohesin loading onto DNA, and methods to conditionally inactivate the ATPase do not yet exist. An arginine finger mutation in the cohesin ATPase substantially slows down cohesin loading onto chromosome but, given enough time to allow for loading, does not compromise sister chromatid cohesion establishment^7^. This observation suggests that the cohesin ATPase is not limiting for cohesion establishment, consistent with the possibility that replisomes pass through inert cohesin rings. Future investigations will also examine how replisome passage through the cohesin ring integrates with others pathways^15,16^ by which cohesin might build cohesive links between sister chromatids.

## Methods

### Protein purification and labelling

#### CMG

A *S. cerevisiae* strain overexpressing the 11 CMG subunits (Y6375, strain genotype details can be found in Extended Data Table 1) was grown overnight at 30°C in YPA (YP with additional 4 mg/L adenine) + 2% raffinose. Cdc45 contained an internal insertion into an unstructured loop, following amino acid 197, of two FLAG epitopes^8^, plus the 12 amino acid S6 sequence (GDSLSWLLRLLN). The next day, the pre-culture was expanded to 12 L at an optical density OD_600_ = 0.5. When OD_600_ reached ∼1.0, protein expression was induced by the addition of 2% galactose (v/v) for 6 h at 30°C. Cells were harvested, washed with water and resuspended in CMG buffer (25 mM HEPES-NaOH pH 7.5, 300 mM KCl, 2 mM Mg(OAc)_2_, 0.02% Tween-20, 0.5 mM TCEP, 10% glycerol) supplemented with protease inhibitors (SigmaFAST, Sigma-Aldrich). This cell suspension was frozen dropwise in liquid nitrogen, crushed by freezer milling and the cell powder was stored at −80°C.

The cell powder was thawed and resuspended in CMG buffer supplemented with protease inhibitors, 0.5 mM AEBSF, and 3000 Units benzonase for 12 L starting culture (Sigma-Aldrich). After stirring for 30 min at 4°C, cell debris was removed by ultracentrifugation in a Ti45 rotor at 40,000 rpm for 60 min. The clarified lysate was transferred onto anti-FLAG M2 agarose beads (Sigma-Aldrich, 4 mL of beads for 12L of culture) equilibrated in CMG buffer and incubated on a rocking platform at 4°C for 4 h. Flag beads were transferred into two 20 ml disposable columns (BioRad), washed with 50 column volumes (cv) CMG buffer and then incubated with ATP wash buffer (CMG buffer with added 1 mM ATP) for 10 min. After further 15 cv CMG buffer wash, proteins were eluted by first incubating for 40 min in 2 cv of CMG buffer supplemented with 0.5 mg/ml 3xFLAG peptide, followed by 20 min in 1 cv of CMG buffer supplemented with 0.25 mg/ml 3xFLAG peptide. The eluted fractions were combined, supplemented with 2 mM CaCl_2_, and incubated overnight at 4°C with calmodulin affinity resin (2 mL resin for 12L of culture). The resin was collected in a 20 mL disposable column, washed with 50 cv calmodulin wash buffer (CMG buffer with 2 mM CaCl_2_), then proteins were eluted with calmodulin elution buffer (CMG buffer with 2 mM EDTA and 2 mM EGTA). The calmodulin eluate was loaded onto a MonoQ PC 1.6/5 column (Cytiva) and separated with a 300-600 mM KCl gradient over 10 cv. Peak fractions were pooled, snap frozen in liquid nitrogen and stored at −80°C.

For fluorophore labelling, CMG in the pooled MonoQ fractions was incubated with phosphopantetheinyl transferase^43^ and LD655-CoA (Lumidyne Technologies, 1:2:5 molar ratio) in the presence of 10 mM MgCl_2_ at 23°C for 1 hour. The labelling reaction was centrifuged for 10 min at 13,000 rpm in a benchtop centrifuge at 4°C followed by removal of phosphopantetheinyl transferase and excess dye using Superose 6 Increase 3.2/300 (Cytiva) chromatography in CMG buffer. Peak fractions were pooled, snap frozen in liquid nitrogen and stored at −80 °C.

#### Cohesin-SNAP

A tetramer *S. cerevisiae* cohesin-SNAP complex was expressed and purified as previously described^27^. The SNAP-tag was fused to the C-terminus of Smc3. Following the heparin chromatography step, the pooled peak fractions were incubated with benzylguanine-LD555 or benzylguanine-LD655 (Lumidyne Technologies) in a protein-to-dye ratio of 1:2 in the presence of 1 mM DTT overnight at 4°C. Excess dye was removed by gel filtration on a Superose 6 Increase 10/300 GL column (Cytiva) equilibrated in gel filtration buffer (50 mM Tris-HCl pH 7.5, 150 mM NaCl, 0.5 mM TCEP, 10% glycerol). The peak fractions were pooled, snap frozen in liquid nitrogen and stored at −80°C.

#### Scc2-Scc4 and Chl1

The *S. cerevisiae* cohesin loader Scc2-Scc4 complex and Chl1 were expressed and purified as previously described^27,35^.

#### Protocatechuate 3,4-Dioxygenase (PCD)

PCD from *Pseudomonas putida* was expressed and purified as previously described with minor changes^44^. T7 express lysY cells (New England Biolabs) carrying a PCD expression vector (pVP91A-pcaHG, Addgene) were grown at 37°C overnight in LB broth supplemented with carbenicillin. In the following morning, cells were diluted 1:100 into 2 x 1L medium and further incubated at 37°C until an OD_600_ of 0.5 was reached. Cells were transferred to 17°C, and once an OD_600_ of 0.7 was reached, the expression of PCD was induced by addition of 0.5 mM IPTG and 10 mg/L Fe(NH_4_)_2_(SO_4_)_2_. The cells were maintained at 17°C for ∼18 h. Next, cells were harvested, resuspended in PCD buffer (50 mM Tris-HCl pH 7.5, 500 mM NaCl, 10 mM Imidazole, 10% glycerol) supplemented with protease inhibitors and flash frozen in liquid nitrogen. Cells were lysed by subjecting them to three consecutive freeze-thawing cycles followed by sonication. Cell debris was removed by centrifugation, and the clarified lysate was applied to a HisTrap (Cytiva) column that had been equilibrated in PCD buffer. The column was washed with PCD buffer containing 20 mM imidazole, followed by elution with two consecutive 15 cv steps of increasing imidazole concentrations (125 mM and 250 mM imidazole, respectively). The peak fractions were pooled and concentrated using centrifugal filters with a 10 kDa molecular weight cut-off (Merck). Finally, the concentrated protein was applied to a Superdex 200 Increase 10/300 size exclusion column (Cytiva) equilibrated in buffer containing 50 mM Tris-HCl pH 7.5, 100 mM NaCl, 0.1 mM EDTA, 10% glycerol. Peak fractions were pooled, flash frozen in liquid nitrogen and stored at −80°C. The activity of purified PCD was measured using an enzymatic activity assay, whereby the decrease in protocatechuic acid (Sigma-Aldrich) was monitored over time as absorbance decrease at 290 nm.

#### Replication proteins

Additional DNA replication factors, Mcm10, Tof1-Csm3, Mrc1, RPA, PCNA, RFC, Ctf4, Pol α, Pol δ, and Pol ε were expressed and purified as previously described^8,9,45^.

#### V5-antibody functionalization

V5-antibody (Bio-Rad) was functionalised with a 20 kDa biotin-PEG-SVA linker (Laysan Bio) through NHS-ester coupling. Biotin-PEG-SVA was dissolved in DMSO to a final concentration of 3 mM. The V5-antibody and the dissolved PEG were mixed in a 1:20 ratio and incubated for two hours at 22°C and agitated at 400 rpm. The reaction was quenched by the addition of 2 µl of 1 M Tris-HCl pH 7.5. The reaction was applied to a Superose 6 Increase 10/300 GL size exclusion column which had been equilibrated in PBS. Three peaks were observed during elution and, to ascertain the number of PEG labels per antibody, peak fractions were analysed by mass photometry (Extended Data Fig. 3a). The single-PEG labelled antibody fractions were pooled, snap frozen in liquid nitrogen and stored at −80°C.

### *In vitro* ensemble assays

#### Cohesin ATPase and DNA loading assays

The rate of ATP hydrolysis by cohesin and its ability to load onto DNA in a salt-resistant manner were determined using previously established assays^4,27^.

#### CMG helicase assay

CMG unwinding was determined following a previously published protocol with minor changes^31^. The forked DNA substrate was assembled by annealing 50duplex-lag-FAM and 50duplex-lead in TE (oligonucleotide sequences can be found in Extended Data Table 2). The mix was heated to 95°C and slowly cooled to 20°C. The assembled fork was purified by TBE gel electrophoresis, electroelution and dialysis against storage buffer (20 mM Tris-HCl pH 8.0, 20 mM NaCl). The fluorescent forked DNA concentration was determined using the fluorescently labelled oligonucleotide as the standard.

For CMG unwinding assays, 10 nM CMG and 3 nM forked DNA were mixed in CMG buffer (25 mM Tris-HCl pH 7.5, 10 mM MgAc_2_, 250 mM potassium glutamate, 0.0025 % Tween 20, 1 mM TCEP, 0.1 mg/mL BSA) on ice. The reaction was started by addition of 3 mM ATP and incubated at 30°C. Aliquots were retrieved at indicated times and quenched by addition to 3xStop buffer (30 mM Tris-HCl pH 7.5, 60 mM EDTA, 1.5% SDS, 15% Sucrose) and separated on a 10% acrylamide TBE gel. The gel was imaged using a fluorescence imager and the unwinding activity was quantified as the amount of unwound 50duplex-lag-FAM using ImageJ.

### Linear forked DNA substrate for single molecule experiments

The linear forked DNA substrate was prepared following previously published protocols with minor changes^30,46^. First, pUber was digest with BstXI (2.5 Units per µg of DNA, New England Biolabs) in NEBuffer 3.1 for 8 hours at 37°C. The reaction was quenched by addition of EDTA (12 mM final) and NaCl (300 mM final), and digested DNA was separated by size exclusion chromatography over a home-made Sepharose CL-4B chromatography column equilibrated in eluent buffer (10 mM Tris-HCl pH 8.0, 12 mM EDTA, 300 mM NaCl). Peak fractions were pooled and dialysed two times for 1 hour against TE (10 mM Tris-HCl pH 8.0, 1 mM EDTA). Capping fragments were generated by annealing oligonucleotides in two separate reactions in 1xTE, blockLd-biteg and blockLg were annealed in a 1:6 molar ratio, 99Lg-biteg, 160Ld and Fork-primer were annealed in a 1:6:60 molar ratio. Annealing reactions were heated to 95°C and, after 2 minutes, the temperature was reduced 1°C every 40 sec until reaching 20°C. The annealed capping fragments were now ligated to linearised pUber using T4 DNA ligase (100 Units per µg of DNA, New England Biolabs) in 1xCutSmart supplemented with 1 mM ATP and 1 mM DTT. The ligation reaction was incubated overnight at 16°C and then quenched with 12 mM EDTA and 30 mM NaCl. Excess capping oligos and T4 DNA ligase were removed by size exclusion chromatography over a Sepharose CL-4B chromatography column equilibrated in eluent buffer. Peak fractions were pooled, aliquoted and snap frozen in liquid nitrogen.

### Flow cell preparation

#### Functionalisation of coverslips

Coverslips used for single-molecule experiments were cleaned, silanised and PEG functionalised based on previous protocols with adjustments^47–49^. Coverslips (24x60 mm, Marienfeld Superior) were placed into a polypropylene staining jar, filled with anhydrous ethanol and sonicated for ∼20 min. The ethanol was removed, and coverslips washed 10 times with purified water. The jar was filled with 5 M KOH and again sonicated for ∼20 min. KOH was removed, and coverslips washed 10 times with purified water. The ethanol and KOH washes were repeated. Coverslips were carefully dried with nitrogen gas and then plasma cleaned for 3 min at max intensity in a PDC-002-CE device (Harrick Plasma). For silanisation, coverslips were first washed several times with acetone, followed by incubation in 2% 3-aminopropyltriethoxysilane (Alfa Aesar) in acetone with shaking. The silane solution was decanted, the coverslips washed once with acetone and then sonicated for 30 sec in acetone. Subsequently, coverslips were washed 20 times with purified water and immediately dried with nitrogen gas.

For 6 coverslips, 75 mg mPEG-succinimidyl valerate (SVA) and 3 mg biotin-PEG-SVA (MW 5000, Laysan Bio Inc.) were mixed and dissolved in 500 µl 0.1 M NaHCO_3_ Three of the silanised coverslips were placed face-up in an empty freezer box, humidified by water at the edges. 150 µl of PEG solution was added and the three remaining coverslips placed on top, face-down, sandwiching the PEG solution. After ∼3 h, the PEGylated coverslips were rinsed, placed into a clean jar and washed 20 times with purified water. Coverslips were dried with nitrogen gas and the PEGylation process was repeated, this time incubating overnight. Rinsed and dried PEGylated coverslips were stored under vacuum for up to 2 weeks.

#### Flow cell assembly

A standard flow cell comprised 7 flow channels, cut into parafilm and sandwiched between one PEGylated coverslip and a quartz coverglass (cleaned in acetone and sonication for 30 min in ethanol). The coverglass contained 7 corresponding pairs of drilled holes. The assembly was secured by melting the parafilm on a heating block for 3 min at 110°C, then sealing the edges with epoxy glue. PE60 tubing (0.76/1.22 mm, Stoelting) was fit into the coverglass holes and sealed with epoxy glue. For side flow experiments, a flow channel comprised one straight channel with a second channel branching at a right angle from the flow cell centre to a second outlet.

### Single-molecule assays

All buffers used in single-molecule experiments were filtered and degassed for at least 30 min immediately before use.

#### DNA tethering

Linear forked DNA was tethered to a PEGylated coverslip surface via biotin-streptavidin interaction. An assembled flow cell was placed onto the microscope, with the outlet tubing of one flow channel connected to a microfluidics pump (Harvard Apparatus), and the inlet placed into sample tubes containing the indicated solutions. Washing and DNA tethering steps were performed at a flow rate of 100 µl/min. First, the channel was flushed with 100 µl blocking buffer BB (20 mM Tris-HCl pH 7.5, 50 mM NaCl, 2 mM EDTA, 0.005%Tween-20) including 0.2 ml/ml NeutrAvidin (Thermo Fisher) and incubated for 30 min without flow. Then the flow was restarted and the channel washed with 300 µl BB. Now 450 µl of 10 pM linear forked DNA in BB, supplemented with 0.2 µM chloroquine (Sigma-Aldrich) was introduced, and untethered DNA was immediately flushed out with further BB.

#### Cohesin loading

Prior to cohesin loading, the surface of the DNA containing flow channel was additionally passivated by incubating for 20 min with 1 mg/ml Ultrapure BSA (Thermo Fisher) in cohesin loading buffer CLB (35 mM Tris-HCl pH 7.5, 25 mM NaCl, 25 mM KCl, 1 mM MgCl_2_, 1 mM DTT, 0.003% Tween-20, 5% glycerol). For cohesin loading, 1.5 nM fluorescently labelled cohesin and 3 nM Scc-Scc4 were introduced into the flow channel at 20 µl/min for 150 µl in CLB supplemented with 1 mM ATP (Jena Bioscience) and 1 mg/ml BSA. Excess and non-topologically loaded cohesin were immediately removed by washing with CMG buffer (25 mM Tris-HCl pH 7.5, 250 mM Potassium glutamate, 10 mM Mg(OAc)_2_, 0.0025% Tween-20, 1 mM DTT) supplemented with 150 mM NaCl, followed by CMG buffer. Cohesin was imaged in CMG buffer supplemented with 150 nM SYTOX Orange or 5 nM YOYO-1 (both Thermo Fisher) and an oxygen scavenging system (see below). Where indicated, the restriction enzyme XhoI (40 units/ml) in CMG buffer supplemented with 150 nM SYTOX Orange, 0.1 mg/ml BSA and oxygen scavenger was introduced into the flow channel at 50 µl/min over 900 µl.

#### CMG loading and translocation

Following BSA passivation as above, 1 nM CMG-LD655 and 10 nM Mcm10 were introduced into the flow channel at 20 µl/min for 100 µl in CLB supplemented with 0.33 mM ATP and 1 mg/ml BSA. Excess CMG was immediately removed by washing with CMG buffer supplemented with 150 mM NaCl, followed by CMG buffer. CMG was activated and translocation was imaged in CMG buffer supplemented with 10 nM Mcm10, 3 mM ATP, 1 mg/ml BSA and oxygen scavenger. Where indicated, CMG was loaded in presence of 0.33 mM ATPγS and 300 nM RPA was included during activation and imaging.

#### CMG-cohesin encounters

Cohesin-LD555 was loaded onto tethered DNA as above. Next, CMG-LD655 was loaded, excess CMG removed, and CMG activated as described. Cohesin-CMG encounters were imaged for 40 min in CMG buffer supplemented with 10 nM Mcm10, 3 mM ATP, 0.1 mg/ml BSA and oxygen scavenger. For experiments with cohesion establishment factors, 30 nM Tof1-Csm3-Mrc1 or Chl1-Ctf4 were added during activation and imaging. The same experimental design was used when imaging converging CMGs.

For experiments with cohesin immobilisation, following loading, cohesin was initially imaged for 5 minutes to identify topologically loaded and freely diffusing complexes. Now, 25 nM V5 PEG-biotin antibody in CMG buffer with 0.1 mg/ml BSA was introduced over 100 µl at 20 µl/min. After incubation for 3 min, the channel was washed with CMG buffer in preparation for CMG loading and activation. For cohesin elution, the flow channel was flushed with CMG buffer supplemented with 50 µg/ml V5 peptide, 0.1 mg/ml BSA and oxygen scavenger, and imaged for a further 25-30 min. The DNA was afterwards stained with 150 nM SYTOX Orange in CMG buffer.

#### Force calculations

Forces generated by CMG or replisome-cohesin encounters that led to DNA stretching prior to CMG or replisome jump events were calculated using a force-extension formula for worm-like chain polymers^38^. DNA tension in the DNA segment in front of CMG (*F*) was calculated using the worm-like chain equation:

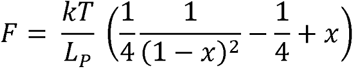

Where *k* is the Boltzmann constant, *T* the absolute temperature, *L_P_* = 50 nm is the persistence length of DNA, and x is the extension ratio of the DNA, which was obtained as follows:

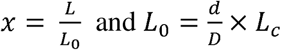

Here, *L* represents the end-to-end distance of the stretched DNA segment immediately before the jump, and *L_0_* is the contour length of that DNA segment. The contour length was calculated assuming that the jump releases and equilibrates tension in the DNA. Therefore, the contour length of the segment is the ratio between the end-to-end distances of the segment *d* and the total DNA *D*, multiplied by the total DNA contour length. For the DNA used here, the contour length of the total DNA was *L_c_* = 6.22 µm.

The DNA length *D* was obtained using the DNA finder tool in Mars^50^. The length of the stretched DNA segment before and after the CMG or replisome jumps were determined by first tracking the CMG using TrackMate, followed by combining the CMG track with the DNA location in Mars. This allowed us to determine where on the DNA CMG was before (*L*) and after the jump (*d*).

#### DNA replication

For additional passivation, the flow channel was incubated with 2% Tween-20 in BB for 10 min prior to DNA tethering. After DNA attachment and equilibration in CMG buffer, 10 nM unlabelled CMG in CMG buffer supplemented with 0.33 mM AMP-PNP and 0.1 mg/ml BSA was flushed into the flow channel at 20 µl/min and incubated for 10 min. Replication was initiated by introducing 60 nM Mcm10, 20 nM Ctf4, 30 nM Tof1-Csm3-Mrc1, 200 nM RPA, 20 nM PCNA, 20 nM RFC, 20 nM Pol α, 20 nM Pol δ and 20 nM Pol ε in CMG buffer (containing 200 mM potassium glutamate) supplemented with 150 nM SYTOX Orange, 125 µM dNTPs, 250 µM NTPs (both Jena Bioscience), 5 mM ATP and 0.1 mg/ml BSA.

#### Replisome-cohesin encounters

Replisome-cohesin encounter experiments were carried out following the same workflow as CMG-cohesin encounters with minor adjustments. The surface was passivated with 2% Tween-20 prior to DNA attachment, and all buffers were supplemented with 0.1 mg/ml BSA. Cohesin-LD655 was loaded, washed and imaged as described. Unlabelled CMG was then loaded, and replication was initiated as detailed above, with the additional inclusion of an oxygen scavenging system. For experiments with immobilised cohesin, the immobilisation and V5 peptide elution steps followed the same procedure as described under CMG-cohesin encounters. For experiments that included LD655-labelled CMG, 1 nM CMG introduced and incubated for 5 minutes in the presence of AMP-PNP, before replication proteins and nucleotides were added.

#### Side flow experiments

The protein and buffer conditions in side flow experiments remained unchanged, unless specified otherwise. The side flow channels were prepared as described above, with the main and side outlets connected to separate microfluidics pumps. First, NeutrAvidin in BB was flushed through both channels and incubated for 30 min. Both channels were washed with 200 µl BB and linear forked DNA (20 pM in BB containing 0.2 µM chloroquine) was introduced through the main channel at 50 µl/min over 250 µl. The unbound DNA was washed sequentially through the main and side channels with BB. After incubating both channels with 1 mg/ml BSA in CLB, cohesin was loaded at 20 µl/min over 150 µl through the main channel. After sequential washes of both channels with CMG buffer containing 150 mM NaCl, cohesin was imaged in CMG buffer supplemented with 5 nM YOYO-1, oxygen scavenger, and 0.1 mg/ml BSA. Now, side flow was applied at 50 µl/min and cohesin immobilised with 50 nM V5 PEG-biotin antibody in CMG buffer with 0.1 mg/ml BSA for 6 min. Both channels were washed with CMG buffer and CMG was loaded at 20 µl/min for 100 µl through the main channel. Excess CMG was removed by washing through both channels using CMG buffer containing 150 mM NaCl, followed by CMG buffer. Finally, CMG was activated, and cohesin-CMG encounters were imaged in CMG buffer supplemented with 10 nM Mcm10, 5 nM YOYO-1, 3 mM ATP, 1 mg/ml BSA and oxygen scavenger while maintaining a constant flow of 5 µl/min through the main channel. After 40 min, cohesin was eluted with 50 µg/ml V5-peptide in CMG buffer supplemented with 5 nM YOYO-1, 0.1 mg/ml BSA and oxygen scavenger.

#### Oxygen scavenging systems

Two oxygen scavenging systems were used to increase fluorescence dye stability. The first consisted of PCD (see above), Protocatechuic acid (PCA) and 6-hydroxy-2,5,7,8-tetramethylchroman-2-carboxylic acid (Trolox) (both Sigma-Aldrich), and was used for imaging of cohesin, CMG and cohesin-CMG encounters. CMG imaging buffer was supplemented with 20 nM PCD, 2.5 mM PCA (from a 650 mM stock in DMSO) and 1 mM Trolox (from a 4 mM stock in CMG buffer). The second oxygen scavenging system comprised glucose oxidase, catalase and glucose and was used for cohesin-replisome encounter experiments. 0.2 mg/ml glucose oxidase (from a 20 mg/ml stock in 50 mM Tris-HCl pH 7.5, 50 mM NaCl and 50% glycerol), 0.035 mg/ml catalase (from a 3.5 mg/ml stock in the same buffer) and 0.5% glucose were added to CMG imaging buffer.

### Data acquisition, processing and analysis

#### Imaging

All experiments were performed on a Nikon Eclipse Ti2-E inverted TIRF microscope equipped with a SR HP Apo TIRF 100x/1.49 oil immersion objective and a LU-NV-D laser bed. The imaging temperature was maintained at 30°C in an electrically heated chamber (Okolab). Cohesin diffusion and photobleaching kinetics were captured every 100 ms for 1-2 min. Other experiments were typically imaged every 20 sec in an area covering 4x4 field of views. The fluorescence signal was spectrally separated using a T635lpxr-UF2 beam splitter (Chroma) and recorded on two Prime 95B sCMOS cameras (Teledyne photometrics) with 0.11 µm x 0.11 µm pixel size. The system was controlled using Nikon NIS-Elements software.

#### Image processing

Fields of view were first separated into individual files and saved in tiff format. Subsequent image processing and analysis were carried out in Fiji (Version 1.54f). Each image was corrected for stage drift using a custom script^30^. Channels were aligned manually using an overexposed SYTOX Orange stain image as a guide. Individual events occurring on DNA were cropped and saved as individual files for later analysis. Kymographs across a 3 pixels wide line were created using the KymoResliceWide plugin (https://doi.org/10.5281/zenodo.4281086).

#### Tracking of CMG translocation and replication

Single fluorescent spots on DNA were tracked using the Dog detector and Linear-motion LAP tracker in TrackMate (v7.13.2)^51^ and the resulting trajectories were exported to Mars^50^. Kinetic change point analysis was used to fit individual rate segments^52^ and the obtained rates were exported for further statistical analysis using Matlab (v.2023b) and GraphPad Prism (v.10.3.0). CMG translocation rates were weighted by the duration of a segment.

#### Tracking of cohesin for photobleaching and MSD analyses

Single fluorescent cohesin on DNA was tracked, and trajectories exported to Mars, as above. The mean intensity of cohesin was plotted over time and kinetic change point analysis was used to fit photobleaching steps^52^. Mean square displacement (MSD) analysis was performed using the Matlab class msdanalyzer^53^. The diffusion coefficient of individual cohesin molecules was estimated from the slope of a linear fit to the first five time delays (0.1-0.5 s). Only coefficients with a goodness of fit of R^2^ < 0.8 were considered.

### Quantification and statistical analysis

Plots showing individual data points were generated using GraphPad Prism (v.10.3.0). The number of molecules n is indicated in each figure, or its legend. All experiments were conducted in at least two biological replicates. Statistical significance was evaluated using an unpaired t-test with statistical significance defined as *p* < 0.05. All error bars are defined in the figure legends and typically represent the standard error of the mean.

## Acknowledgements

We thank J. Lewis and L. Spenkelink for guidance, C. Bouchoux, G. Pobegalov, and H. Yardimci for reagents and advice, the Crick Chemical Biology Science Technology Platform for dye conjugation, as well as D. Ramirez Montero and our laboratory members for discussion and critical reading of the manuscript. This work was supported by Wellcome Trust Investigator Awards (219527/Z/19/Z to J.F.X.D. and 220244/Z/20/Z to F.U.) and The Francis Crick Institute, which receives its core funding from Cancer Research UK, the UK Medical Research Council, and the Wellcome Trust (cc2125 to M.I.M.; cc2002 to J.F.X.D; cc2137 to F.U.)

## Author contributions

S.G., J.F.X.D. and F.U. conceived the study, S.G. performed all experiments, M.I.M. advised on forces and their calculation, S.G. and F.U. wrote the manuscript with input from all coauthors.

---

**Extended Data Fig. 1.**
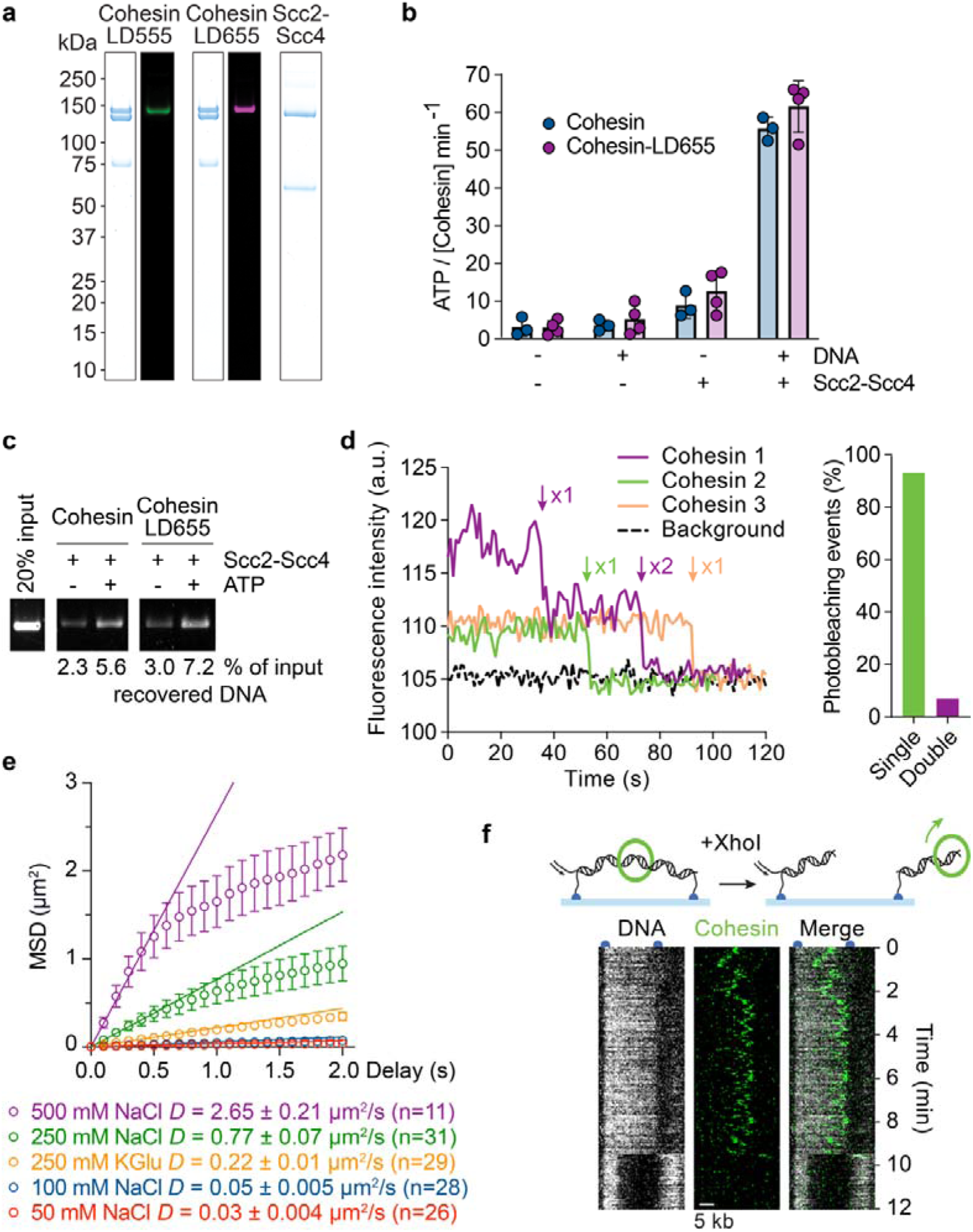
Ensemble and single molecule analysis of budding yeast cohesin. **a,** Purified and fluorophore-labelled budding yeast cohesin tetramer complexes (Smc1, Smc3, Scc1 and Scc3) were analysed by SDS-polyacrylamide gel electrophoresis and Coomassie blue staining or in-gel fluorescence. The purified, unlabelled Scc2-Scc4 cohesin loader complex was analysed alongside. **b,** ATP hydrolysis rates of unlabelled and LD655-labelled cohesin in presence or absence of DNA and Scc2-Scc4. Each point shows the result from independent biological replicate experiments. Bars show the mean, error bars the standard deviation. **c,** Gel image of DNA recovered in a cohesin loading assay with unlabelled and LD655-labelled cohesin, in the presence Scc2-Scc4, with or without added ATP. The amount of recovered DNA was quantified, compared to the input. **d,** Example photobleaching traces of cohesin on DNA, with single and double bleaching steps highlighted by arrows. The percentage of single and double step events is shown (n = 44). **e,** Mean square displacement (MSD) as a function of delay time for cohesin on DNA imaged at increasing salt concentration. The mean and standard error are plotted for each delay time. Data points up to delay times of 0.5 seconds were fitted with linear regressions, where the slope denotes the diffusion coefficients (*D*, indicated). **e,** Schematic and representative kymograph of cohesin sliding off the free end of surface-tethered DNA following restriction digestion.

**Extended Data Fig. 2.**
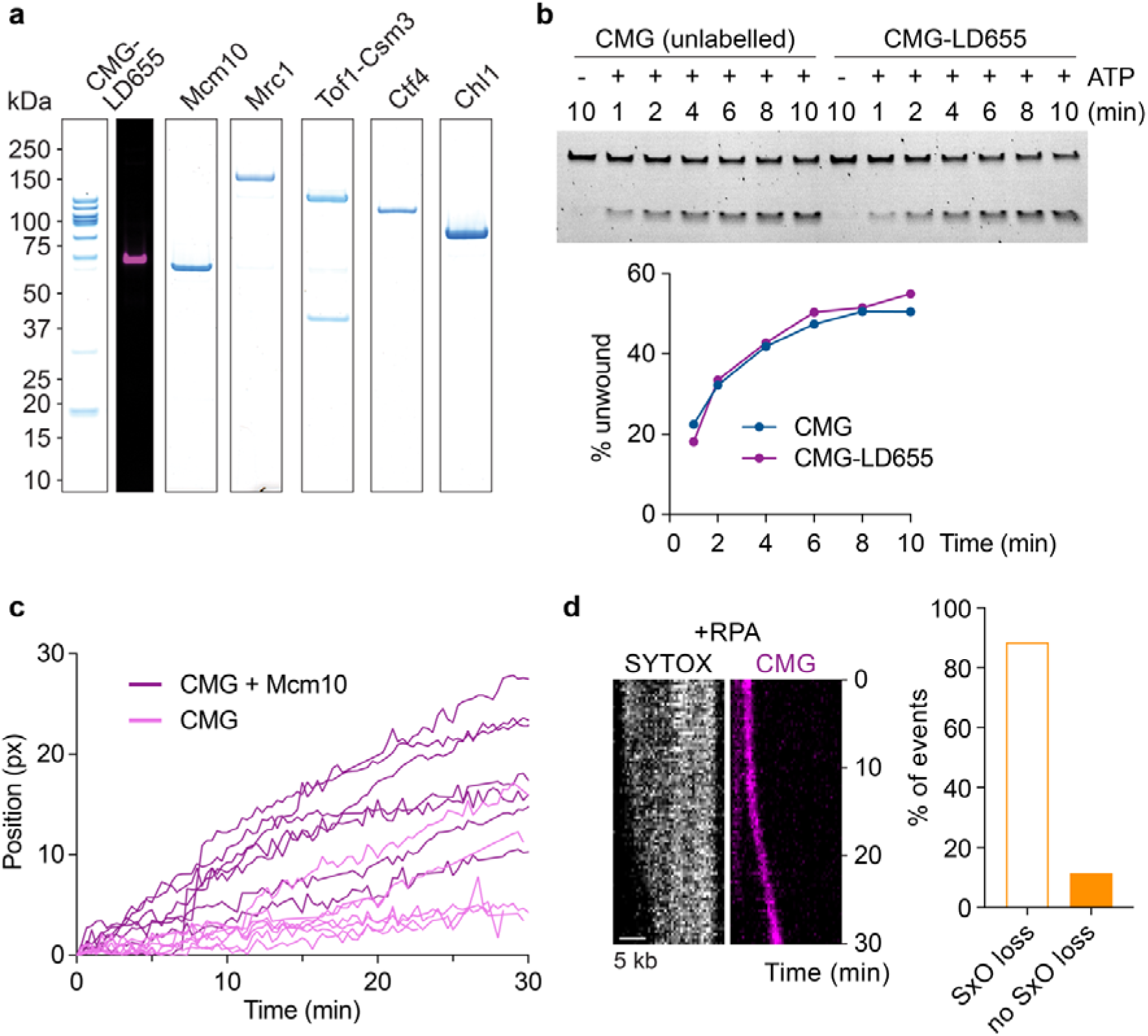
Ensemble and single molecule analysis of budding yeast CMG. **a,** Purified and fluorophore-labelled budding yeast CMG was analysed by SDS-polyacrylamide gel electrophoresis and Coomassie blue staining or in-gel fluorescence. Purified, unlabelled Mcm10, Mrc1, Tof1-Csm3, Ctf4 and Chl1 were analysed alongside. **b,** CMG unwinding assay comparing unlabelled and labelled CMG. The fluorescence gel image shows the fork substate and unwound single stranded product. Quantification of the fraction of unwound DNA over time is shown underneath. **c,** Example trajectories of CMG positions along the DNA as a function of time, in the presence or absence of Mcm10. **d,** Representative kymograph of DNA unwinding by CMG in the presence of RPA. SYTOX Orange preferentially stains dsDNA, with loss of the SYTOX Orange signal indicating DNA unwinding. The percentage of CMG translocation events that were accompanied by SYTOX Orange signal loss is indicated (n = 190).

**Extended Data Fig. 3.**
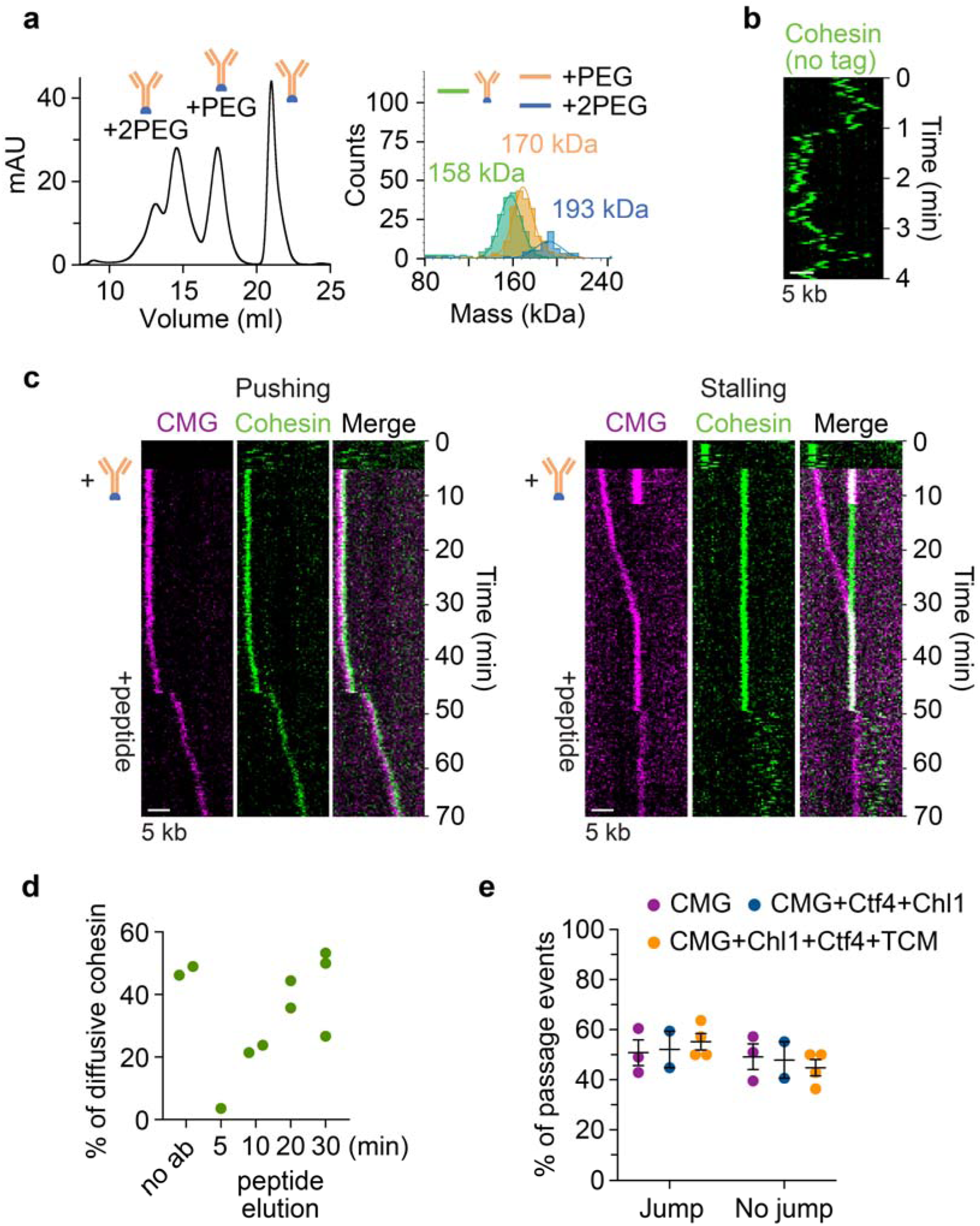
CMG passing immobilised cohesin. **a,** Size exclusion chromatography and corresponding mass photometry profiles of functionalised V5 antibody after reacting with 20 kDa (∼150 nm) biotin-PEG-succinimidyl valerate. Molecular weights correspond to unreacted, single-, and double-labelled antibody, respectively. Fractions from the single labelled peak were pooled. **b,** Representative kymograph of DNA-bound cohesin after introducing V5 antibody lacking the biotin-PEG tag. **c,** Representative kymographs of CMG pushing a previously immobilised cohesin (left), and of CMG stalling at immobilised cohesin (right). **d,** Percentage of diffusive cohesin observed before antibody immobilisation, and at the indicated times after peptide addition. Each value represents the fraction of diffusive over static cohesin in one field of view, obtained in biological replicate experiments. Static cohesins are mostly surface-rather than DNA-bound molecules. **e,** Classification of CMG passage events with or without discernible jump, in the absence or presence of the indicated cohesion establishment factors. Each data point is the result of a biological replicate. The black lines represent the mean and standard error.

**Extended Data Fig. 4.**
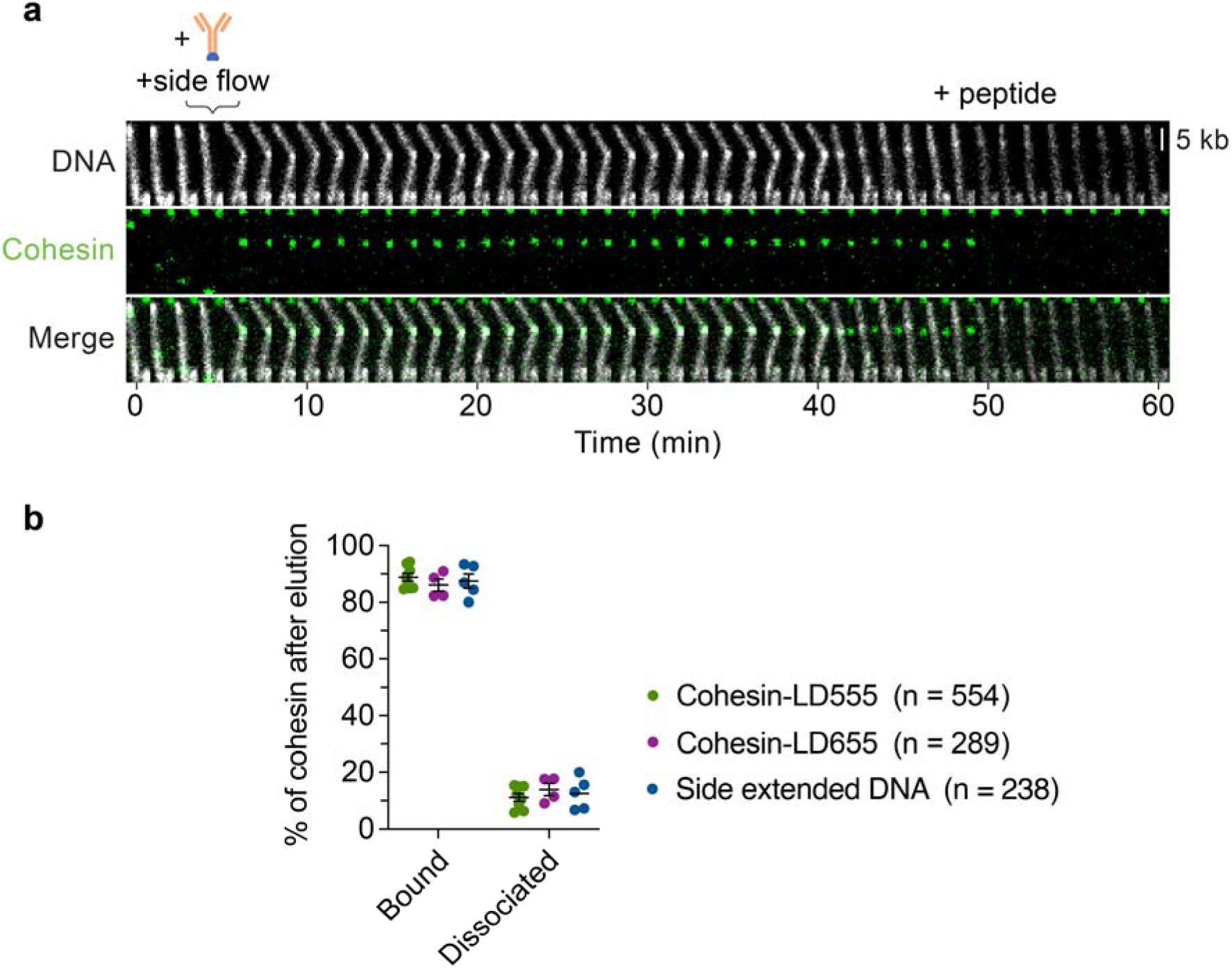
Cohesin anchoring and release of side flow-stretched DNA. **a,** Representative montage of side flow-stretched DNA, tethered by cohesin after the side flow was stopped. Cohesin released the DNA at ∼40 minutes, presumably due to spontaneous unloading. Upon V5 peptide addition, cohesin was eluted from the surface (at ∼50 minutes). **b,** Quantification of cohesin stability on flow-stretched DNA. Cohesin was immobilised following loading onto DNA without side flow (Cohesin-LD555 and Cohesin-LD655), or with side flow (Side extended DNA). Following peptide elution, we counted the fraction of cohesins that remained DNA-bound or that dissociated. The analysis revealed that, irrespective of DNA geometry, around 10% of cohesins spontaneously unloaded from the DNA during these procedures.

**Extended Data Fig. 5.**
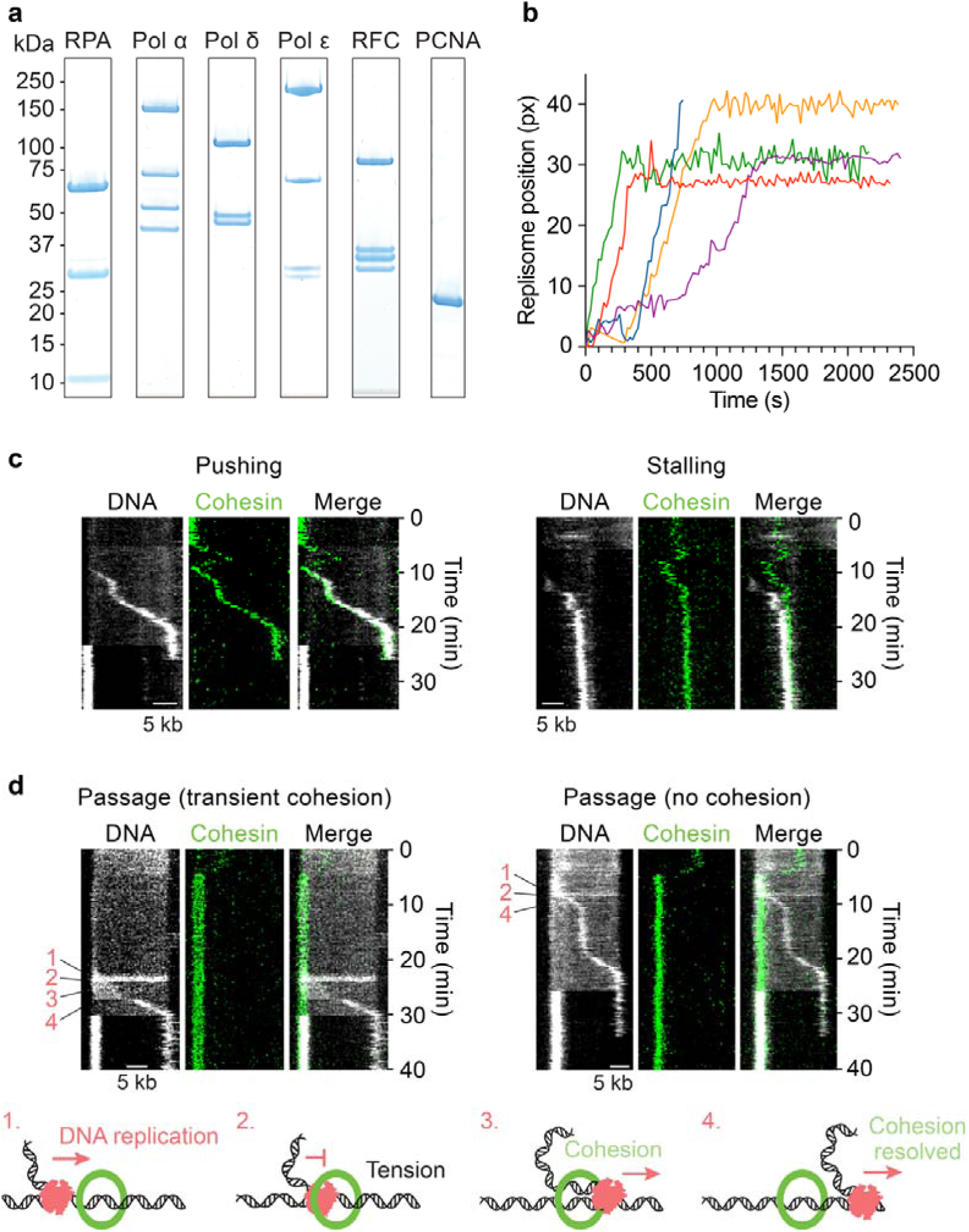
Single molecule replisome cohesin encounters. **a,** Purified budding yeast RPA, Pol α, Pol δ, Pol ε, RFC and PCNA were analysed by SDS-polyacrylamide gel electrophoresis and Coomassie blue staining. **b,** Representative DNA replication trajectories, tracking the centre of mass of the leading strand replication product over time. **c,** Representative kymographs of replisome-cohesin encounters that resulted in cohesin pushing (left) or in replisome stalling (right). **d,** Representative kymographs of replisome-cohesin encounters that resulted in cohesin passage with only transient cohesion establishment (left) or without cohesion establishment (right). The schematics illustrate how the free leading strand DNA end might slip through cohesin rings.

**Extended Data Fig. 6.**
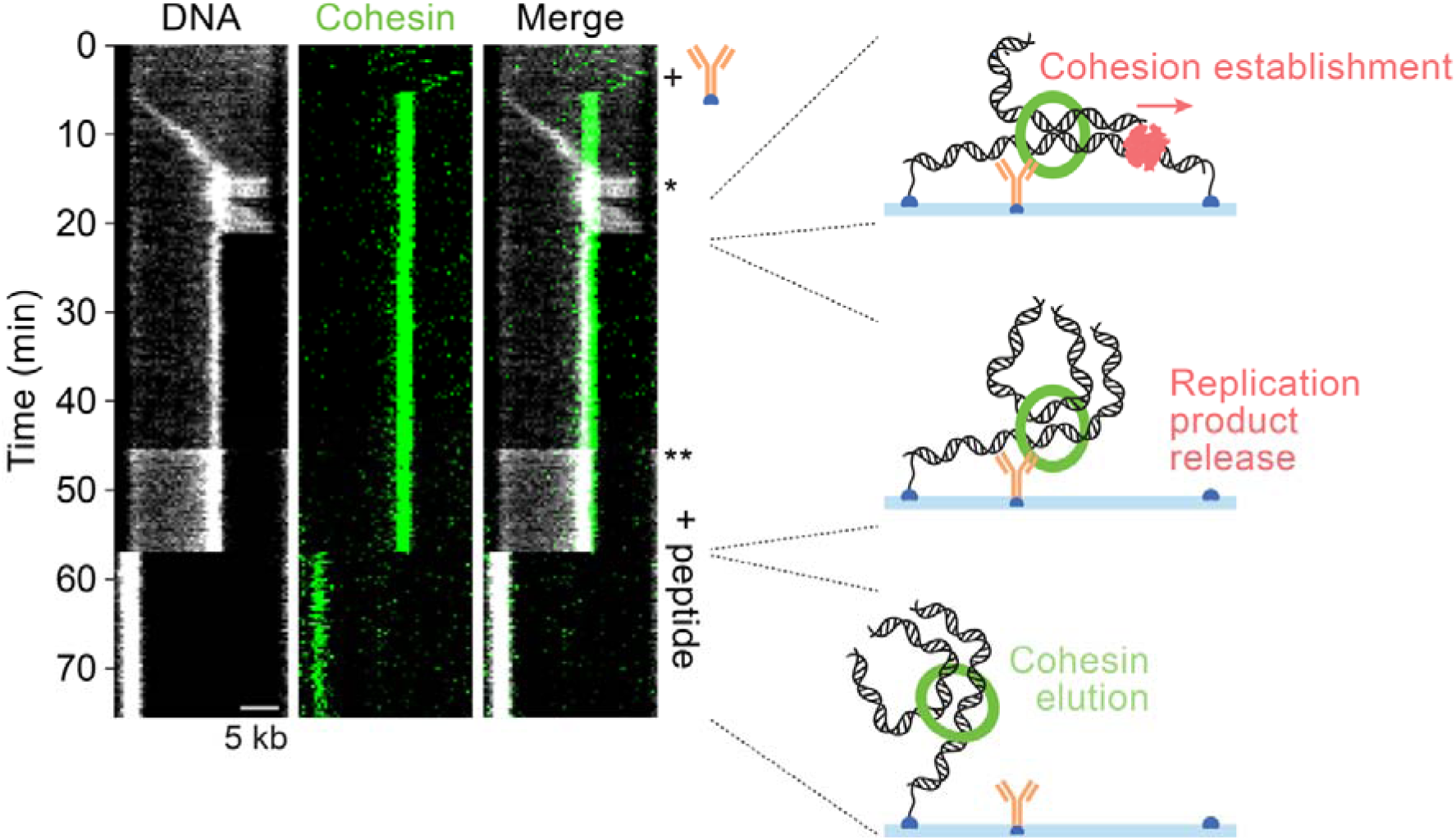
Replisome encounter with immobilised cohesin. Representative kymograph of a replisome encounter with immobilised cohesin, with schematics that illustrate sequential events. Upon initial encounter, DNA tension builds up ahead of the replisome (*). Once the replisome passes cohesin, cohesion has been established and the two DNA replication products continue to elongate. Upon completion of DNA replication, the replication products are released from the flow cell surface (synthesis of the final nucleotides displaces the DNA anchor from the coverslip) but remain entrapped by cohesin. After V5 peptide addition (**, when fresh SYTOX Orange in the elution buffer results in a DNA signal gain) cohesin eventually elutes and the DNA snaps back to its upstream coverslip anchor.

